# Novel KIF22 Variants Disrupt Mitosis in Human Chondrocytes and Expand SEMDJL2 Mechanisms

**DOI:** 10.64898/2026.03.11.711192

**Authors:** Amila Šemić, Kazette Yuen Yu Chan, Pricila Bernardi, Karina C. Silveira, Cynthia Silveira, Denise P. Cavalcanti, Peter Kannu, Jason Stumpff

**Affiliations:** Department of Molecular Physiology and Biophysics, University of Vermont, Burlington, VT 05405; Skeletal Dysplasia Group, Medical Genetics Department, Science Medical Faculty, University of Campinas (UNICAMP), Brazil; Pediatrics Department of University Hospital, Federal University of Santa Catarina, Brazil; Department of Medical Genetics, University of Calgary. Canada; Neurology Department, University of Campinas (UNICAMP), Campinas, SP, Brazil

## Abstract

Spondyloepimetaphyseal dysplasia with joint laxity, type 2 (SEMDJL2) is a rare skeletal disorder caused by pathogenic variants in KIF22, a mitotic chromokinesin that generates polar ejection forces (PEF) to ensure proper chromosome alignment and segregation. Although prior work showed that SEMDJL2-associated KIF22 hotspot variants impair chromosome segregation in epithelial cells, how these variants affect chondrocyte mitosis remains incompletely understood. Here, we analyzed the effects of the hotspot variant R149Q, a recently reported recessive variant R49Q, and two newly identified heterozygous variants, P144T and E222Q, in human chondrocytes. Both novel variants were identified in individuals with classic SEMDJL2 features. P144T and E222Q retained PEF-generating activity, whereas R49Q displayed reduced PEFs, consistent with their respective inheritance patterns. Live cell imaging revealed that all variants disrupted mitosis. The heterozygous variants (P144T, E222Q, R149Q) dominantly impeded anaphase chromosome segregation and spindle pole separation, supporting reclassification of P144T and E222Q as likely pathogenic. In contrast, R49Q caused milder, partially penetrant segregation defects, consistent with reduced and dysregulated motor activity. Together, our results define two mechanistic classes of KIF22 dysregulation: constitutive activation in heterozygous variants, which fail to down-regulate KIF22 at anaphase onset, and mixed-state dysregulation in the recessive R49Q variant, which exhibits partial loss of polar ejection force activity coupled with incomplete inactivation during anaphase. These findings broaden the mechanistic framework for how KIF22 variants perturb mitosis in chondrocytes and expand the genotypic landscape associated with SEMDJL2.

## INTRODUCTION

Spondyloepimetaphyseal dysplasia with joint laxity, type 2 (SEMDJL2; MIM 603546), is a rare skeletal dysplasia characterized by short stature, midface hypoplasia, joint laxity with multiple dislocations, especially of the knees, progressive scoliosis, and a typical radiological pattern featuring remarkable delay of carpal maturation with slender metacarpal/phalanges (leptodactyly), small/fragmented epiphysis, metaphyseal vertical striation on knees and ankles, and characteristic slender and tapered femoral neck. (Hall et al., 1998; Kim et al., 2009; Nishimura et al., 2003; Rossi et al., 2005; Boyden et al., 2011; Min et al., 2011; Dubail et al., 2024). Genetic variants in the mitotic motor protein KIF22 have been identified as pathogenic for SEMDJL2 (Boyden et al., 2011; Min et al., 2011; Thompson et al., 2022). However, the molecular pathological mechanism linking KIF22 (MIM 603213) dysfunction to skeletal abnormalities remains unclear.

KIF22, also known as kinesin-like DNA-binding protein (Kid), is a plus-end directed chromokinesin that binds to both microtubules and chromosomes (Levesque and Compton, 2001; Tokai et al., 1996). As a motor protein, KIF22 generates pushing forces on chromosome arms, called polar ejection forces (PEF), that facilitate chromosome alignment and compaction (Brouhard and Hunt, 2005; Levesque and Compton, 2001, Stumpff et al., 2012). Upon mitosis onset, KIF22 actively generates polar ejection forces, which push chromosome arms away from the spindle poles and toward the metaphase plate at the center of the spindle (Levesque and Compton, 2001). At the metaphase to anaphase transition, KIF22 is inactivated, reducing polar ejection forces on chromosomes and allowing them to separate and move toward the spindle poles (Su et al. 2016; Soeda et al. 2016). This activity is known to be regulated by phosphorylation; KIF22 is activated by phosphorylation at Thr463 at the start of prophase by Cdk-1/Cyclin-B and dephosphorylated and inactivated at the metaphase-to-anaphase transition to turn off its activity (Oshugi et al., 2003; Soeda et al., 2016).

Thompson et al. (2022) functionally characterized SEMDJL2 hotspot variants (affecting residues P148 and R149) in KIF22 and found that expression of KIF22 variants disrupts chromosome segregation and spindle elongation during anaphase. The consequences of these mitotic defects in HeLa and RPE-1 cells include cytokinesis failure, abnormal daughter cell nuclear morphology, and reduced proliferation. The KIF22 variants still were able to generate polar ejection forces, similar to those of KIF22-WT, indicating a retention of motor function. Additionally, expression of phospho-mimetic KIF22 constructs (T463D) to prevent inactivation of polar ejection force generation at the metaphase to anaphase transition phenocopied the effects of expressing SEMDJL2 KIF22 variants (Thompson et al. 2022). This was consistent with previous studies where live oocytes expressing KIF22-T463D constructs underwent chromosome recongression (Soeda et al. 2016). Taken together, these data suggest that KIF22 SEMDJL2 variants continue to generate PEF after the metaphase-to-anaphase transition, which impairs chromosome segregation and spindle pole separation in anaphase B.

SEMDJL2-associated KIF22 variants reported to date are dominant, heterozygous missense variants, predominantly located within the α2-region of the motor domain (**Figure 1A**). An exception was reported by Dubail et al. (2024), who described a homozygous motor-domain variant, c.146G>A p.(Arg49Gln), in 3 affected individuals, along with unaffected heterozygous carriers. Individuals homozygous for the R49Q variant presented the same SEMDJL2 phenotype as previously reported. The R49 residue is located near the beginning of the motor head domain of KIF22 and is highly conserved across human kinesins, and across KIF22 orthologs (Dubail et al., 2024). Functional studies revealed that both dominant and recessive variants preserved KIF22 mRNA and protein levels but impaired aggrecan secretion and extracellular matrix formation, implicating disrupted cartilage biology (Dubail et al., 2024). Whether the R49Q variant affects KIF22’s mitotic functions remains unknown.

**Figure 1:**
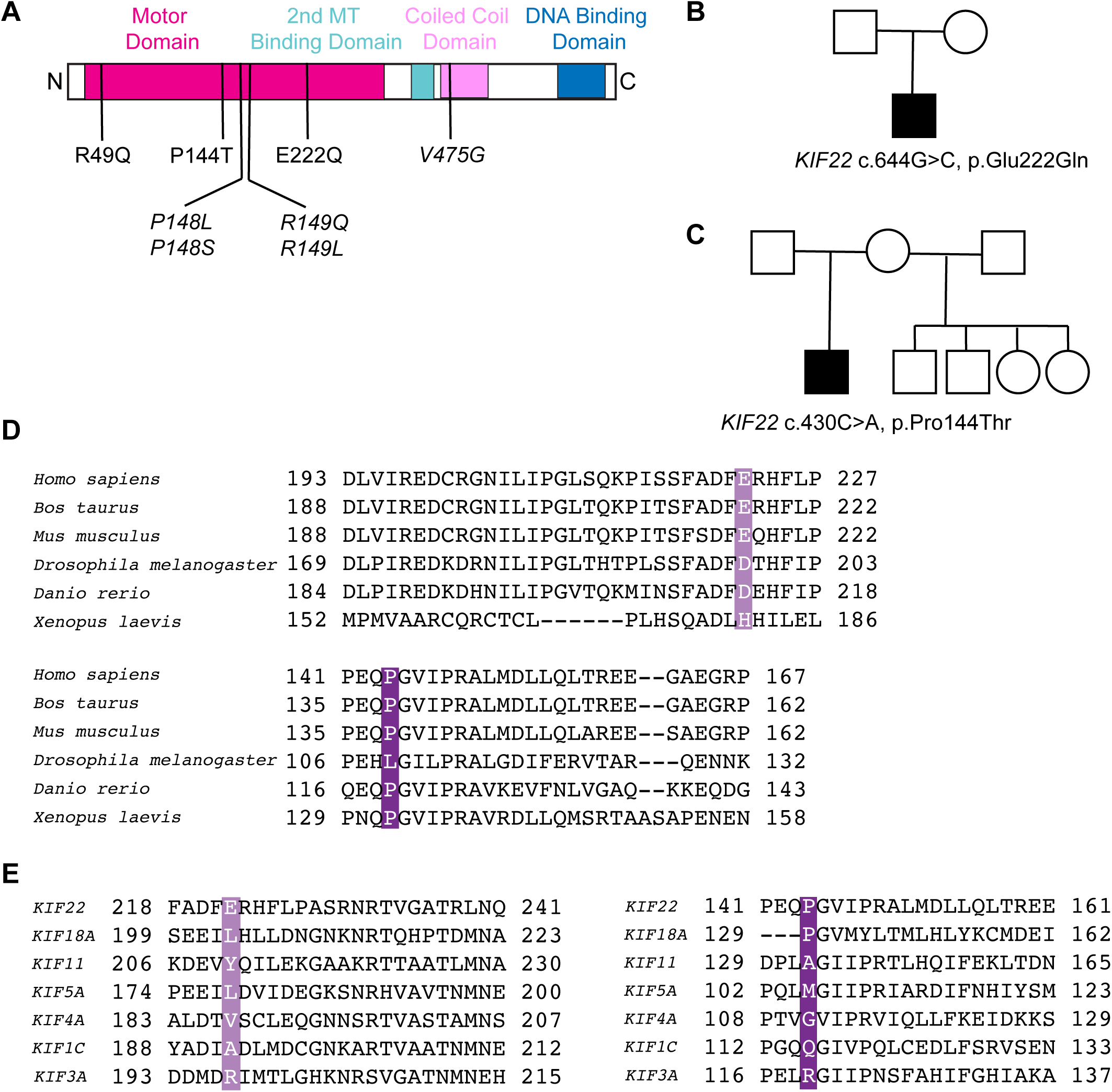
Discovery of SEMDJL2 patients with novel KIF22 variants. **(A)** A linear schematic of the KIF22 protein with functional domains labeled. Locations of novel variants are indicated in the motor domain (R49Q, P144T, E222Q). Locations of previously characterized variants are indicated in italics (hotspot variants *P148L, P148S, R149Q, R149L,* and the tail domain variant *V475G*). **(B)** Pedigree of the patient carrying the p.E222Q variant. **(C)** Pedigree of the patient carrying the p.P144T variant. **(D)** Alignments of amino acid sequences of kinesin-10 family members across species. **(E)** Alignments of amino acid sequences of human kinesin motor domains.

Concurrently, Kawaue et al. (2024) demonstrated that KIF22 is highly expressed in murine chondrocytes of the proliferative zone of the growth plate, consistent with other qRT-PCR analyses in mice (Boyden et al., 2011; Min et al., 2011). Interestingly, mice that were heterozygous for a non-functional KIF22 truncation mutant (amino acids 1-159) had shorter growth plates and long bones compared to wild type mice. Cultured chondrocytes from cartilage of the same mutant mice had a modest increase in abnormal spindle morphology, though the mechanistic cause of these abnormalities was not determined. Furthermore, overexpression of a KIF22-P143L variant (equivalent to P148L in SEMDJL2 patients) in cultured murine chondrocytes led to abnormal mitotic spindle organization, reduced chondrocyte proliferation, and decreased expression of cartilage matrix genes (Kawaue et al., 2024). These results suggest that both loss of function and gain of function variants in KIF22 can negatively impact chondrocyte mitosis. However, the details of these findings conflict somewhat with data indicating that KIF22-P148L does not impact spindle formation and maintenance in epithelial cells (RPE-1 and HeLa) but instead affects spindle pole separation in anaphase (Thompson et al. 2022).

Thus, it remains unclear how KIF22 SEMDJL2 variants affect mitosis in chondrocytes. To address this question and provide a mechanistic examination of mitotic spindle function in human chondrocytes expressing SEMDJL2 KIF22 variants, we analyzed the effects of the hot spot variant R149Q, the homozygous variant R49Q, and two novel heterozygous KIF22 variants identified in patients with clinical presentations that fall within the known SEMDJL2 spectrum (P144T and E222Q). We found that all variants disrupt mitosis by impairing chromosome segregation and spindle pole separation to varying degrees in chondrocytes, consistent with patterns observed in epithelial cells (Thompson et al., 2022). Our functional characterization of P144T and E222Q supports their reclassification from variants of uncertain significance (VUS) to likely pathogenic variants, expanding the documented genotypic spectrum of SEMDJL2-associated *KIF22* variants. By examining both non-hotspot and recessive variants, this work broadens the mechanistic model for how these dysplasia-associated variants dysregulate KIF22 activity during mitosis. Our findings suggest two distinct classes of dysfunction: constitutive activation in the heterozygous variants and mixed-state dysregulation observed in the homozygous variant.

## RESULTS

### Reporting novel KIF22 variants as pathogenic for SEMDJL2

Here, we report two variants of uncertain significance (VUS), *KIF22* c.430C>A, p.(P144T) and *KIF22* c.664G>C, p.(E222Q) (**Figure 1B, 1C**). Variants of uncertain significance (VUS) are genetic variants for which pathogenicity cannot be confidently assigned due to limited population frequency data and the absence of functional validation. Both patients were heterozygous for their respective KIF22 variants. Clinically, both patients demonstrated features consistent with the classic presentation of SEMDJL2, including disproportionate short stature, characteristic epiphyseal and metaphyseal abnormalities, joint laxity, scoliosis, and delayed bone maturation, especially in the carpal region and characteristic slender femoral neck. Although these two amino acid substitutions fall outside the previously described hotspot residues P148 and R149, they are still located within the motor domain of KIF22 (**Figure 1A**), suggesting a high possibility of pathogenicity. P144 and E222 are conserved residues across KIF22 orthologs, but less well conserved among kinesin homologs (**Figure 1D, 1E**). P144 is part of Loop 5 in the motor head domain, which is a structurally conserved loop near the ATP binding site, and E222 is part of α-helix-3.

Proposita 1 is an 18-year-old female who was first evaluated in early childhood for disproportionate short stature and skeletal abnormalities. Radiographic evaluation at 2 years of age in view of short stature demonstrated delayed carpal ossification, metaphyseal changes of the humeri, shortened and bowed femora, and irregularities of the spine, prompting further investigation for an underlying skeletal dysplasia. Clinical examination at the age of 16 revealed disproportionate shortening of the lower body segments. Musculoskeletal examination identified restricted elbow extension, small hands, and significant shortening and bowing of the proximal lower limbs. Her cognitive functioning, hearing, and vision were normal. A comprehensive 358-gene skeletal dysplasia panel (Invitae) was performed, identifying the *de novo* heterozygous missense variant of uncertain significance (VUS): *KIF22* c.430C>A (p.Pro144Thr) (**Figure 1B**). Her clinical phenotype was suggestive of SEMDJL2. Propositus 2 is a Brazilian male who was evaluated at age 6 for short stature and skeletal abnormalities. At 6 years of age, clinical examination revealed disproportionate short stature, with lower-limb shortening (height of 99 cm [<< 3rd centile]) and relative macrocephaly (head circumference of 53 cm [75th centile]), mild craniofacial dysmorphisms (an upturned nose and a depressed nasal bridge), scoliosis, genu valgum, and flat feet with prominent calcaneus. Radiographic evaluation revealed a delay of bone ossification (absence of carpal ossification and of interphalangeal epiphysis), short long bones, small femoral heads and epiphysis in the knees, slender femoral neck, some metaphyseal irregularity with vertical striation in the metaphysis of the distal femur and proximal tibiae. The overall clinical and radiographic findings were consistent with the SEMDJL2 hallmark phenotype. The Sanger sequencing of KIF22 identified a *de novo* heterozygous missense VUS: KIF22 c.664G>C, p.(Glu222Gln). Then, an in-house-designed skeletal dysplasia panel (Silveira et al., 2021), confirmed the same results (**Figure 1C**).

### Motor activity of dysplasia-associated KIF22 variants in chondrocytes

SEMDJL2-associated variants in KIF22 were previously shown to cause mitotic defects in HeLa-Kyoto and RPE-1 cells (**Figure 1**, Thompson et al., 2022), as well as in cultured murine chondrocytes (Kawaue et al., 2024). These mitotic defects primarily included impaired anaphase chromosome segregation and spindle elongation, cytokinesis failure, and abnormal daughter cell nuclear morphology in human epithelial cells (Thompson et al., 2022), abnormal spindle morphology in murine chondrocytes (Kawaue et al., 2024), and reduced proliferation seen in both cell types. To determine how SEMDJL2-associated variants affect KIF22 activity and mitotic spindle function in human chondrocytes, we generated CHON-001 cells that inducibly express GFP-tagged WT KIF22, KIF22-R149Q (hot spot variant), KIF22-P144T, KIF22-E222Q, or KIF22-R49Q (homozygous variant).

At the onset of mitosis, KIF22 is activated by CDK1-cyclin B phosphorylation (Soeda et al., 2016). KIF22 binds both chromosomes on its tail and spindle microtubules on the motor head domain and generates polar ejection forces, which collectively push chromosome arms toward the center of the metaphase plate (Brouhard and Hunt, 2005). This activity is important for chromosome congression and alignment during metaphase, and KIF22 is the main generator of polar ejection forces in cells (Brouhard and Hunt, 2005; Li et al., 2016). These polar ejection forces can be unambiguously measured on monopolar spindles, where KIF22 activity pushes the chromosomes away from the centrosomes at the center of the monopole. When KIF22 is knocked down, chromosome arms collapse toward the center of the monopoles due to reduced polar ejection forces, and this distance is decreased (**Figure 2A, 2B**) (Thompson, Vandal, and Stumpff, 2022; Levesque and Compton, 2001). Previously tested SEMDJL2 KIF22 variants (P148L, P148S, R149L, R149Q, V475G) were able to produce polar ejection forces in cells in this assay similar to that of KIF22-WT and were determined to be functional (Thompson et al., 2022). To evaluate the activity of newly identified KIF22 SEMDJL2 variants, we treated CHON-001 cells expressing KIF22-GFP constructs with monastrol and either control or KIF22 siRNA and then measured the positions of chromosomes as a metric for polar ejection force activity (**Figure 2A**). As expected, we found that KIF22 depletion led to decreased distances between chromosomes and centrosomes in monopolar CHON-001 cells. Chondrocytes expressing only GFP provide polar ejection force measurements for endogenous KIF22, which positioned chromosomes at a mean distance from the poles of 3.0 ± 0.05 µm and this was reduced to 1.2 ± 0.1 µm when KIF22 was knocked down. Inducing expression of KIF22-WT-GFP in cells that also had endogenous KIF22 knocked down restored chromosome distance from the poles to 3.1 ± 0.05 µm. Overexpression of heterozygous variants also restored polar ejection forces to the same levels observed for KIF22-WT (P144T: 2.8 ± 0.8 µm, E222Q: 3.1 ± 0.04 µm) with endogenous KIF22 knockdown. KIF22 KD did not significantly change PEF produced by the heterozygous variants (**Figure 2B**). These data indicate that the novel heterozygous KIF22 variants can move chromosomes to a similar extent as wild-type KIF22. In contrast, we found that the homozygous KIF22 variant produced significantly less polar ejection forces in the absence of endogenous KIF22 compared to wild-type KIF22 (R49Q + KIF22 KD: 2.6 ± 0.6 µm; GFP + Control KD: 3.0 ± 0.05 µm; WT + KIF22 KD: 3.1 ± 0.05 µm). In cells expressing both endogenous KIF22 and KIF22-R49Q, average chromosome distances to the pole were normal (3.0 ± 0.05 µm) and decreased with KIF22 KD (2.6 ± 0.06 µm), indicating that KIF22-R49Q is a loss-of-function allele, consistent with pathological effects only occurring in patients that are homozygous for the variant (**Figure 2B**).

**Figure 2:**
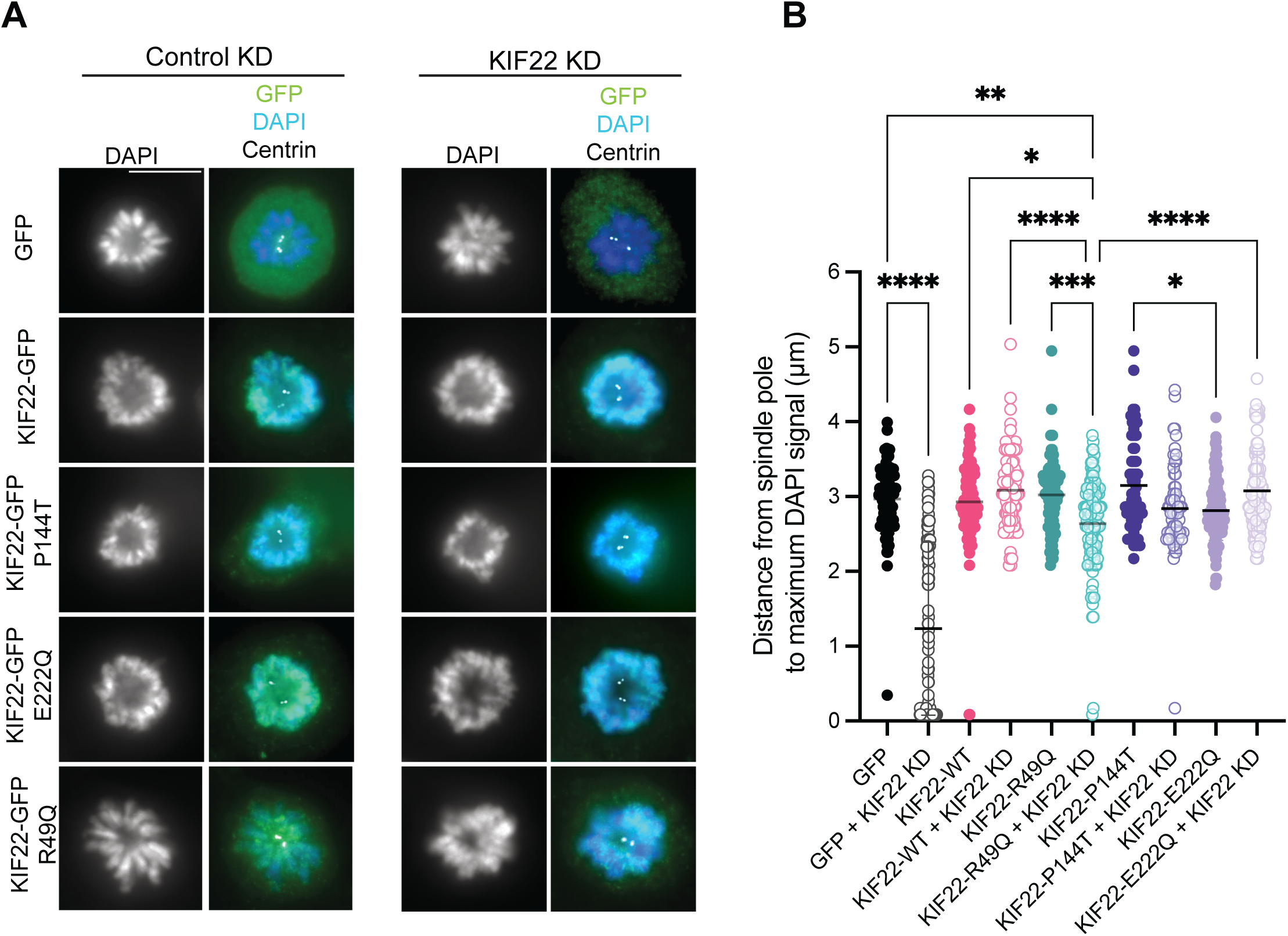
Polar ejection forces produced by SEMDJL2 variants. Relative polar ejection forces (PEF) were assessed by measuring the maximum distance from chromosomes to spindle poles in chondrocytes after knocking down KIF22 and expressing GFP-tagged KIF22 SEMDJL2 variants. 2-3 hours prior to fixation, treatment with monastrol (KIF11 inhibitor) collapsed mitotic spindles to monopoles and arrested cells in mitosis. **(A)** Immunofluorescence images of chondrocyte monopolar spindles. Images are representative of 3 independent experiments. Cells were fixed and stained with DAPI to label DNA, anti-centrin to label centrosomes, and anti-GFP to label GFP or KIF22-GFP constructs. **(B)** Relative PEF measured as the distance from the spindle pole to the maximum DAPI signal intensity, which was calculated using radial plot profiles in ImageJ. Data were compared by running a one-way ANOVA with Šídák’s multiple comparisons test; p-value style: <0.05 (*), <0.01 (**), <0.001 (***), and <0.0001 (****). If no significance is indicated, the result was not significant (>0.05). Scale bar is 10μm.

### Expression of KIF22 variants disrupts anaphase in chondrocytes

To determine how SEMDJL2 variants affect chromosome segregation in chondrocytes, we performed live imaging on dividing chondrocytes expressing WT or SEMDJL2-associated KIF22 variants (R149Q, P144T, E222Q, or R49Q). Expression of GFP-tagged KIF22 variants was induced in CHON-001 cells labeled with SiR-tubulin to observe spindle microtubules (**Figure 3A)**. Anaphase defects were quantified by tracking the distance between the separating chromosome masses using the KIF22-GFP signal and the distance between the two spindle poles using the SiR-tubulin signal in live movies (**Figure 3B-E)**. The average maximum distance between separating chromosome masses during anaphase was 14.8 ± 0.37 µm in cells expressing KIF22-WT (**Figure 3B, 3D**). This distance was reduced to 5.0 ± 0.61 µm and 8.6 ± 0.73 µm in cells expressing KIF22-E222Q and KIF22-P144T respectively (**Figure 3B, 3D).** Spindle elongation during anaphase B was similarly impacted; the average maximum distance between spindle poles during anaphase was 21.3 ± 0.51 µm for cells expressing KIF22-WT and was reduced to 14.4 ± 0.40 µm and 17.4 ± 0.05 µm for cells expressing KIF22-E222Q and KIF22-P144T, respectively (**Figure 3C, 3E**). This effect was dominant, observed in the presence of endogenous KIF22. To compare these effects to those induced by a SEMDJL2 hotspot variant, we analyzed chromosome segregation and anaphase spindle elongation in CHON-001 cells expressing KIF22-R149Q (**Supplemental Movie 3**). As expected from the results in HeLa and RPE-1 cells (Thompson et al., 2022), expression of KIF22-R149Q significantly reduced the maximum distance chromosomes were able to separate to 4.7 ± 0.73 µm (**Figure 3D**), and the maximum anaphase spindle length to 15.8 ± 0.56 µm (**Figure 3E**). Thus, the severity of chromosome segregation defects caused by the R149Q and E222Q variants are comparable, with the P144T phenotype being quantitatively less severe.

**Figure 3:**
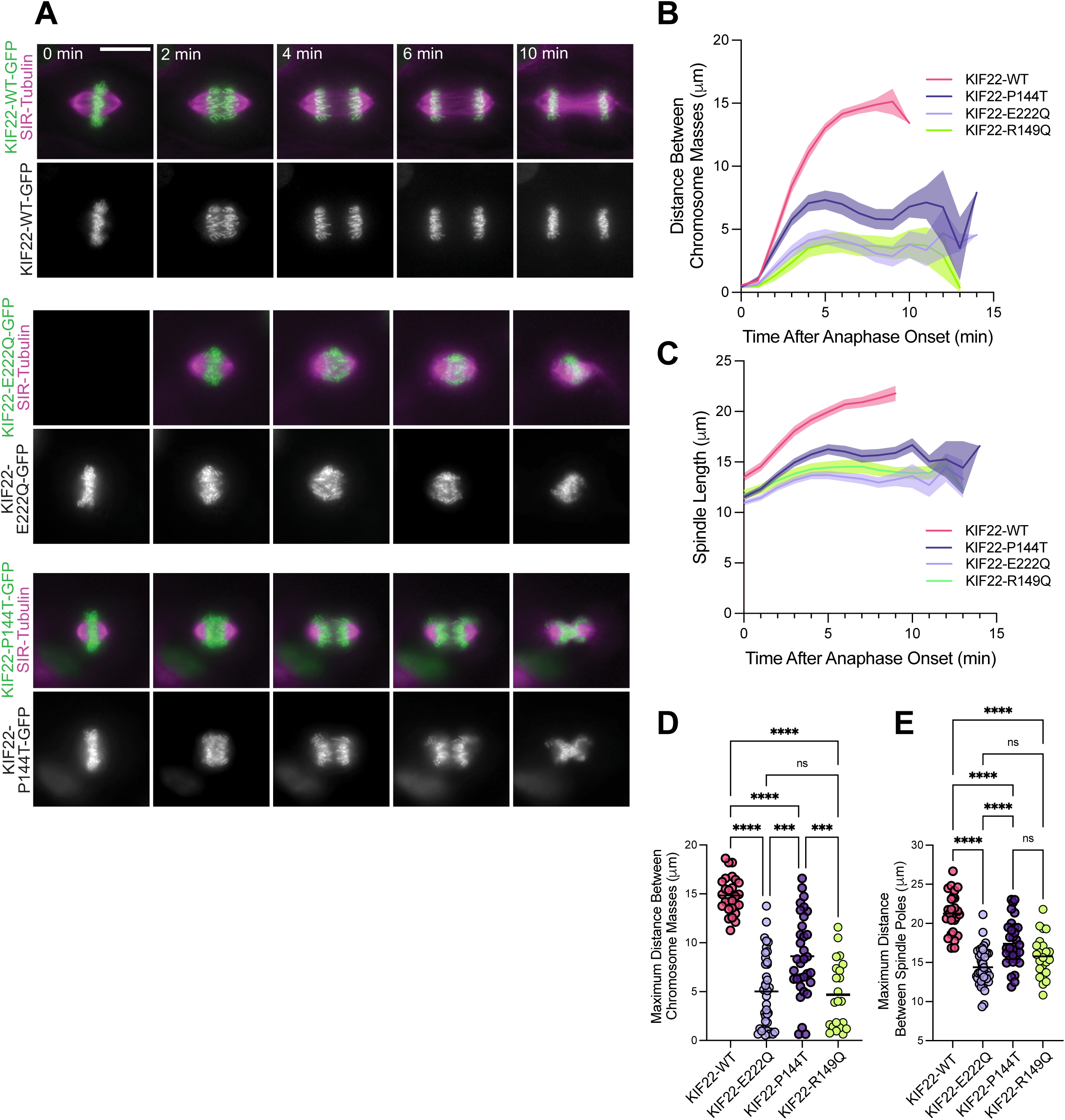
Novel heterozygous SEMDJL2 variants cause anaphase B defects in chondrocytes. To evaluate effects of KIF22 SEMDJL2 variants on mitosis, GFP-tagged KIF22 constructs were expressed in immortalized chondrocytes. **(A)** Timelapse images from live movies in chondrocytes expressing either KIF22-WT-GFP, KIF22-P144T-GFP, or KIF22-E222Q-GFP, or KIF22-R149Q-GFP (see Supplemental Movie 3). Mitotic spindles were labeled with SiR-tubulin (magenta). Movies were captured at 1 frame per minute. Images are representative of at least 3 independent experiments. Scale bar is 15 μm**. (B)** Distance between separating chromosome masses was tracked using the KIF22-GFP signal in chondrocytes. Lines represent the mean and the shaded area denotes SEM. **(C)** Distance between spindle poles after anaphase onset in chondrocytes. Lines represent the mean and the shaded area denotes SEM. **(D)** The maximum distance between separating chromosome masses was calculated for each division. **(E)** The maximum distance between spindle poles was calculated for each division. **Cell counts**: KIF22-WT: 26 cells; KIF22-E222Q: 38 cells; KIF22-P144T: 33 cells; KIF22-R149Q: 23 cells from at least 3 independent experiments per cell line. Data were compared by running a one-way ANOVA with Tukey’s test for multiple comparisons: p-value style: <0.05 (*), <0.01 (**), <0.001 (***), and <0.0001 (****). If no significance is indicated, the result was not significant (>0.05).

To test how the R49Q loss of function variant impacts anaphase in chondrocytes, we analyzed chromosome segregation and spindle elongation in the CHON-KIF22-R49Q cell line following treatment with either control or KIF22 siRNA to simulate a homozygous phenotype **(Figure 4A-E)**. To complement this analysis, we additionally analyzed chromosome segregation in CHON-001 cells expressing a GFP-only construct as a control. If loss of KIF22 activity disrupts anaphase in chondrocytes, then cells expressing KIF22-R49Q in the absence of endogenous motor should behave similarly to GFP expressing cells with KIF22 knocked down. To visualize chromosomes, CHON-GFP cells were labeled with Spy-DNA for live imaging (**Figure 4A**). Surprisingly, we observed that KIF22-R49Q expressing cells with control KD displayed a modest but significant reduction in the average maximum distance between chromosomes during anaphase compared to the GFP-only with control KD (GFP: 14.3 ± 0.37 µm; R49Q: 11.4 ± 0.42 µm) (**Figure 4B, 4D**). The chromosome separation defect caused by KIF22-R49Q expression worsened with KIF22 KD, lowering it to 8.9 ± 0.40 µm (**Figure 4B, 4D**). Conversely, chromosomes separated further in the GFP-only cells with KIF22 KD (increase to 17.2 ± 0.37 µm) (**Figure 4B, 4D**). These data suggest that although it has reduced function compared to KIF22-WT, KIF22-R49Q does impair anaphase chromosome segregation, with a milder phenotype compared to the heterozygous variants. The greater severity of mitotic defects observed with KIF22 knockdown is consistent with patient phenotypes for this variant, in which heterozygous carriers are unaffected, while skeletal developmental abnormalities occur only in individuals homozygous for the KIF22-R49Q variant (Dubail et al., 2024).

**Figure 4:**
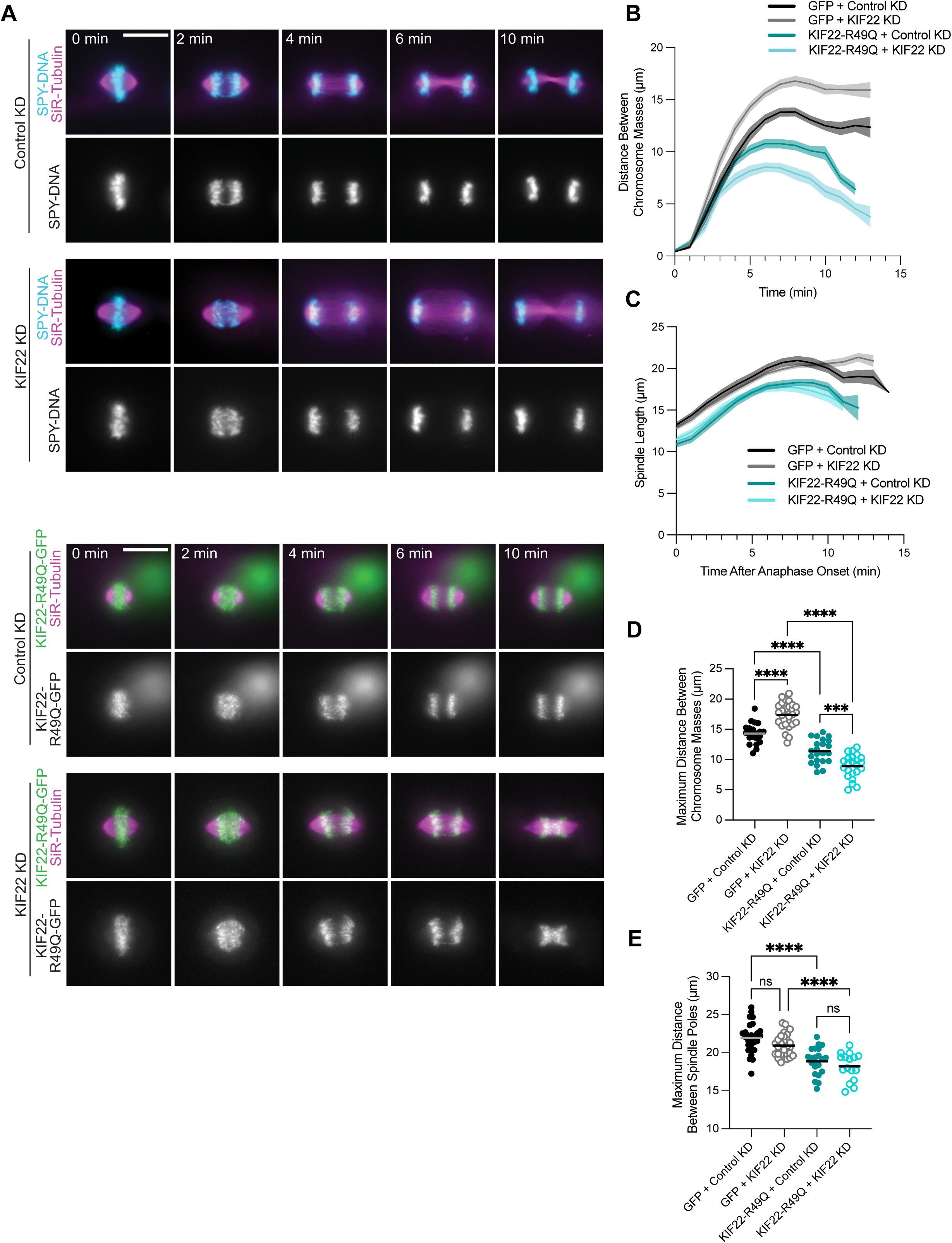
Homozygous SEMDJL2 variant causes mitotic defects in chondrocytes. To evaluate effects of the KIF22-R49Q variant on mitosis, GFP-tagged KIF22 constructs were expressed in immortalized chondrocytes. Cells were induced to express GFP-tagged constructs by treating cells with 2 µg/mL doxycycline. Chondrocytes expressing a GFP-only (null) construct were used as a control. **(A)** Timelapse images from live movies in chondrocytes expressing either GFP or siRNA resistant KIF22-R49Q-GFP and treated with either control or KIF22 siRNA. Mitotic spindles were labeled with SiR-tubulin (magenta), and chromosomes were labeled with Spy-DNA in the GFP line only. Movies were captured at 1 frame per minute. Images are representative of at least 3 independent experiments. Scale bar is 15 µm. **(B)** Distance between separating chromosome masses was tracked using the KIF22-GFP signal in the KIF22-R49Q line or the Spy-DNA signal in the GFP only line. Lines represent the mean and the shaded area denotes SEM. **(C)** Distance between spindle poles after anaphase onset in chondrocytes. Microtubules were labeled with SiR-tubulin. Lines represent the mean and the shaded area denotes SEM. **(D)** The maximum distance between separating chromosome masses was determined for each division. **(E)** The maximum distance between spindle poles was determined for each division. **Cell counts**: KIF22-R49Q (Control KD): 22 cells; KIF22-R49Q (KIF22 KD): 23 cells; GFP (Control KD): 21 cells; GFP (KIF22 KD): 29 cells. Data is from at least 3 independent experiments per cell line and per siRNA condition. Data were compared by running a one-way ANOVA with Tukey’s test for multiple comparisons: p-value style: <0.05 (*), <0.01 (**), <0.001 (***), and <0.0001 (****). If no significance is indicated, the result was not significant (>0.05).

Interestingly, the average maximum spindle length was only modestly reduced in cells expressing KIF22-R49Q with control KD compared to the heterozygous variants (WT: 21.3 ± 0.51 µm; R49Q: 18.9 ± 0.38 µm; P144T: 17.4 ± 0.05 µm; E222Q: 14.4 ± 0.40 µm; R149Q: 15.8 ± 0.56 µm), and KIF22 KD had no effect on spindle elongation for either the GFP or the KIF22-R49Q line compared to their respective control KD condition (**Figure 3E**, **Figure 4E**). The GFP only line displayed a maximum spindle length of 22.0 ± 0.37 µm with control KD and 21.0 ± 0.27 µm with KIF22 KD, while the KIF22-R49Q expressing cells had maximum anaphase spindle lengths of 18.2 ± 0.46 µm with control KD and 18.9 ± 0.38 µm with KIF22 KD (**Figure 4E**). Since spindle pole separation was not as severely affected by the homozygous variant, and it has a reduced capacity to produce PEF (**Figure 2**), the data suggest that KIF22-R49Q perturbs anaphase through a mechanism distinct from the dominant heterozygous variants.

### Novel KIF22 variants disrupt mitosis and cause nuclear morphology defects in daughter cells

Expression of SEMDJL2 variants in epithelial cells leads to lagging chromatin near the cleavage furrow and the formation of lobed nuclei (Thompson et al., 2022). To determine if KIF22-SEMDJL2 variants affect nuclear shape in chondrocytes, we measured nuclear solidity (the ratio of actual nuclear area to the area of a convex shape enclosing it) in cells after inducing expression of GFP-tagged KIF22 variants and treating cells with control siRNA or KIF22 siRNA for 48 hr (**Figure 5A**). A nuclear solidity value of 1 indicates a perfectly round shape. We classified nuclei as abnormal if their solidity value was less than those of the fifth percentile solidity value in GFP-only expressing chondrocytes (solidity value with control siRNA treatment = 0.966). Solidity values were reduced in cells expressing the SEMDJL2 variants (**Figure 5B**), indicating an increase in abnormal nuclear shapes. Chondrocytes were sensitive to overexpression of KIF22-WT, resulting in 19.6% of cells having abnormally shaped nuclei (**Figure 5C)**. This was not surprising, since overexpression of KIF22 could disrupt the balance of forces in mitotic spindles, similarly impairing anaphase (Thompson et al., 2022). Expression of KIF22-P144T increased the percentage of abnormal nuclei to 67.9% (control KD) and 75.6 % (KIF22 KD), while KIF22-E222Q resulted in an increase of 46.7% (control KD) and 54.8% (KIF22 KD) abnormal nuclei (**Figure 5C**). Interestingly, the severity of nuclear morphology defects does not follow trends observed for chromosome segregation defects, since the separation defects were worse with expression of E222Q compared to P144T, while the reverse is true for nuclear morphology. The homozygous variant KIF22-R49Q modestly increased the percentage of abnormal nuclei to 41.9% with control KD. Knockdown of endogenous KIF22 caused a more dramatic increase to 61.2% (**Figure 5C**). We also observed that KIF22 KD in GFP-only expressing chondrocytes caused a small increase in the percentage of cells with abnormal nuclei (Control KD: 4.4 %, KIF22 KD: 14.2%). Together, these data demonstrate that expression of SEMDJL2 variants results in a large increase in abnormally shaped nuclei likely due to instances of impaired chromosome segregation.

**Figure 5:**
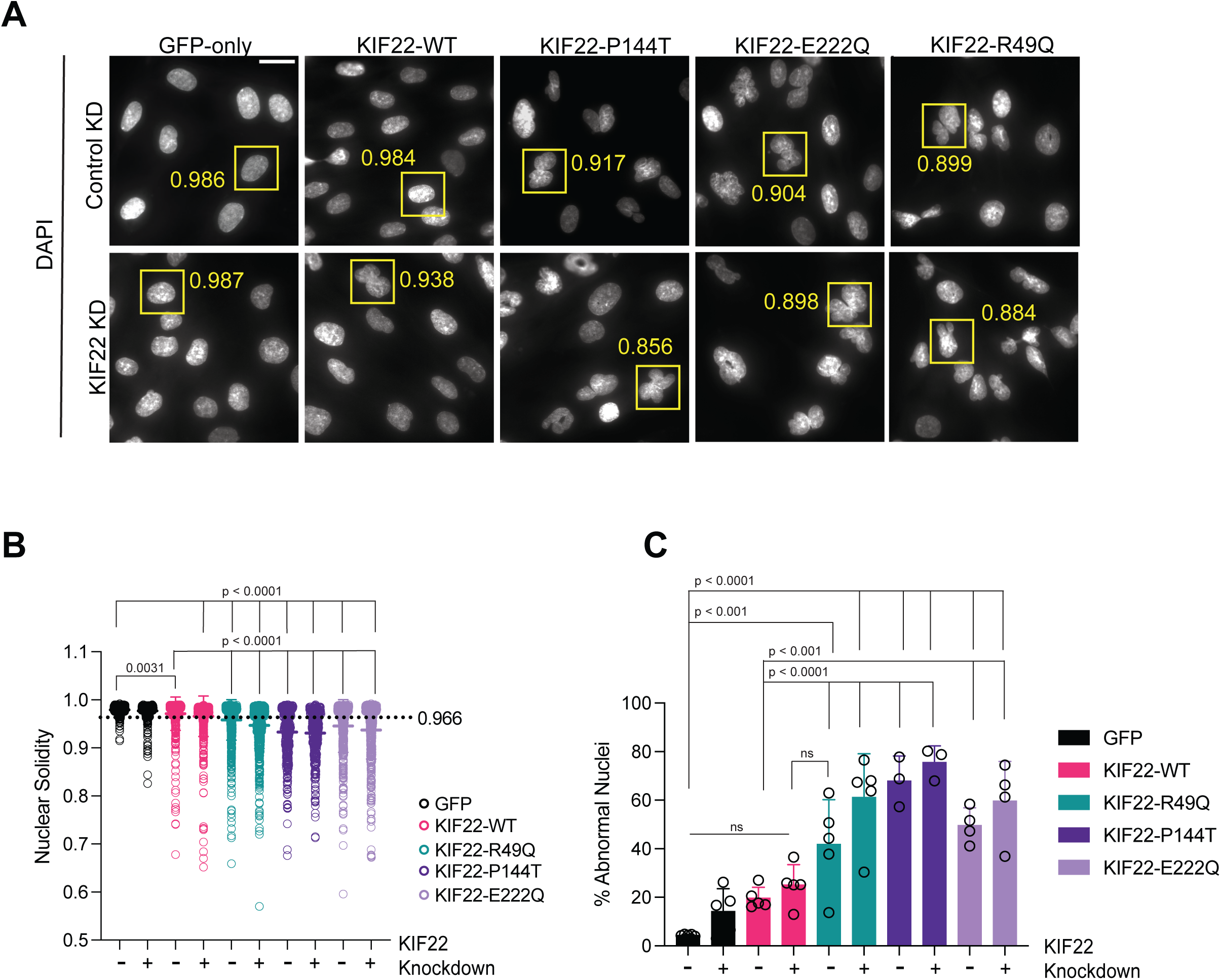
Mitotic defects caused by novel SEMDJL2 variants result in abnormal nuclear morphology. Nuclear morphology was assessed in interphase chondrocytes after 48 hr treatment with control or KIF22 siRNA and 16-18 hr treatment with doxycycline to induce expression of KIF22 constructs. **(A)** Immunofluorescence images of nuclei stained with DAPI from each chondrocyte cell line with control or KIF22 KD. Yellow boxes indicate example nuclei and their solidity values as measured in ImageJ. Scale bar is 20 µm. **(B)** Solidity measurements for chondrocyte nuclei expressing KIF22-GFP constructs. Solidity was measured using ImageJ and a mask of the DAPI signal. Dashed line represents the fifth percentile solidity value of nuclei in control cells (GFP-only cell line with control KD = 0.966). Data were compared by running a one-way ANOVA with Tukey’s test for multiple comparisons. Each condition was compared to GFP control KD or to KIF22-WT control KD. If no significance is indicated, the result was not significant (>0.05). **(C)** Percentage of cells with abnormal nuclear shape, determined by a solidity value less than 0.966. Circles represent average percentage of abnormal cells in one experiment for each condition. Data were compared by running a one-way ANOVA with Šídák’s test for multiple comparisons. Each condition was compared to GFP control KD or to KIF22-WT control KD. If no significance is indicated, the result was not significant (>0.05). Cell counts: **GFP only** – Control KD: 700 cells; KIF22 KD: 666 cells. **KIF22-WT** – Control KD: 716 cells; KIF22 KD: 710 cells. **KIF22-R49Q** – Control KD: 880 cells; KIF22 KD: 716 cells. **KIF22-P144T** – Control KD: 374 cells; KIF22 KD: 361 cells. **KIF22-E222Q** – Control KD: 617 cells; KIF22 KD: 607 cells. Data from at least 3 experiments per cell line.

## DISCUSSION

SEMDJL2 arises from pathogenic disruptions to KIF22, yet the basis by which specific KIF22 variants perturb mitosis and skeletal development has remained incompletely understood. By analyzing both novel heterozygous substitutions and a recently described recessive allele in a physiologically relevant chondrocyte model, this study expands the mechanistic framework for how distinct classes of patient-derived KIF22 variants interfere with mitotic progression. We demonstrate that novel heterozygous KIF22 variants as well as the hotspot variant KIF22-R149Q disrupt anaphase chromosome segregation and spindle elongation in live chondrocyte cells, consistent with the cellular phenotypes demonstrated for the established hot-spot variants in HeLa and RPE-1 cells (Thompson et al., 2022). Interestingly, the individual carrying the E222Q variant presented with a more severe clinical phenotype compared to the patient carrying the P144T variant, which correlated with the results from the in-vitro cellular assays, where expression of the KIF22-E222Q variants caused more severe chromosome segregation defects in anaphase. However, the severity of nuclear morphology defects correlated less well with patient phenotypes. Additionally, in contrast to spindle defects reported by Kawaue et al., (2024) we did not observe abnormal spindle morphology prior to anaphase in any of our chondrocyte cell lines.

A key conceptual insight from our work is that not all dysplasia-associated KIF22 variants destabilize the same aspect of KIF22 regulation. The two newly identified heterozygous variants, KIF22-P144T and KIF22-E222Q, retain the ability to produce polar ejection forces, yet fail to properly down-regulate KIF22 at anaphase onset. This pattern aligns them mechanistically with previously characterized constitutively active SEMDJL2 hotspot and phosphomimetic KIF22 constructs (Thompson et al., 2022; Soeda et al., 2016), suggesting that these substitutions compromise the regulatory mechanism that inactivates KIF22 at the metaphase to anaphase transition. This persistent KIF22 activity during anaphase B erroneously produces polar ejection forces that impede spindle pole separation and interfere with chromosome segregation, offering a mechanistic explanation for how these variants disrupt mitotic fidelity (**Figure 6**).

**Figure 6:**
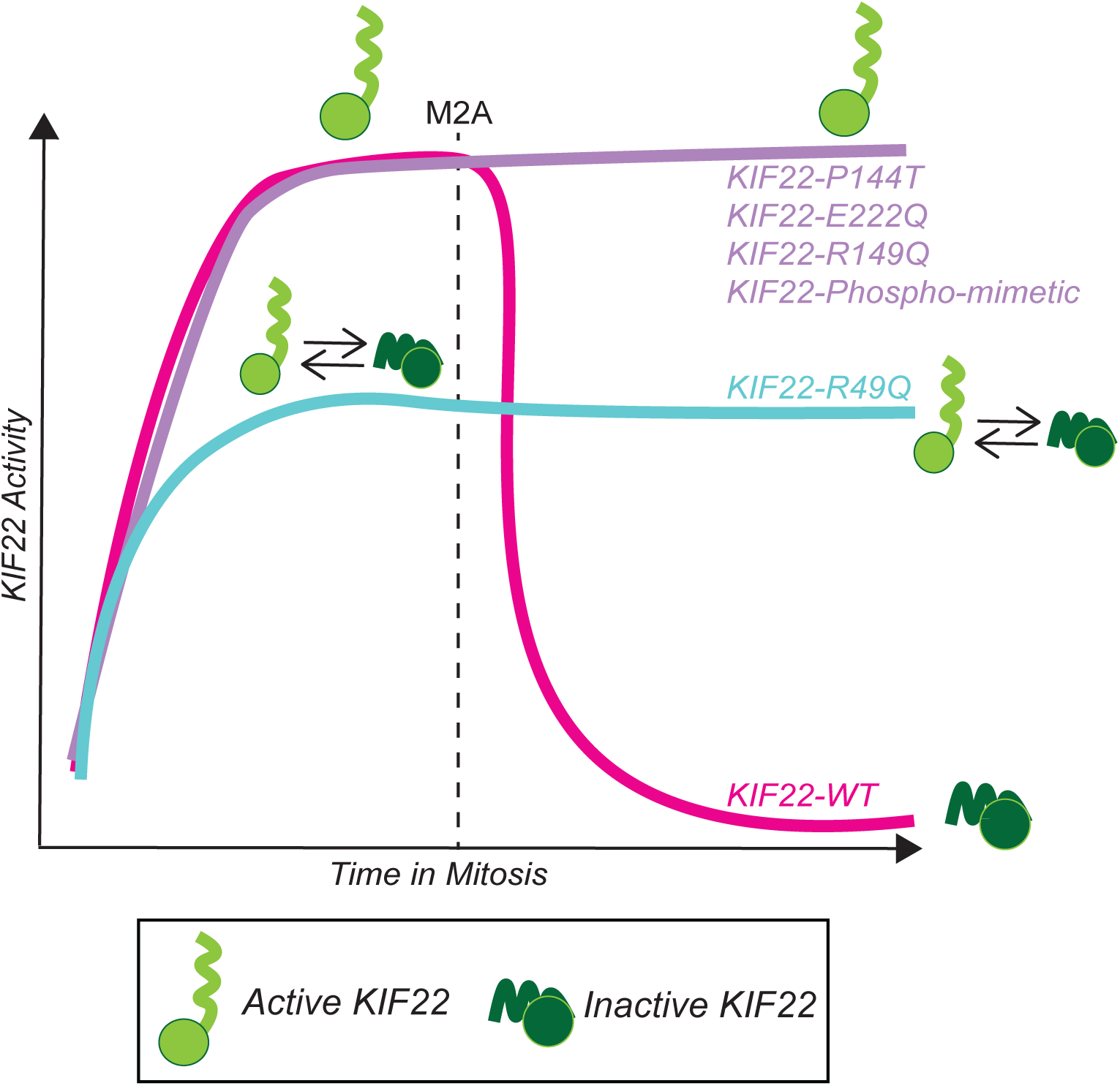
Model of KIF22 activity throughout mitosis with SEMDJL2 variants. KIF22-WT activity is increased via phosphorylation at mitosis onset and is decreased via dephosphorylation at the metaphase-to-anaphase transition (M2A). Heterozygous KIF22 SEMDJL2 variants continue to produce PEF into anaphase, disrupting chromosome segregation. The homozygous R49Q variant cannot regulate its activity during mitosis, leading to a reduced level of PEF before and after M2A. The mechanism of KIF22 inactivation is unknown but depicted here as a head-to-tail autoinhibition as a possible example.

The recessive R49Q variant presents another regulatory scenario. Rather than behaving as a fully active or inactive allele, R49Q appears to compromise both KIF22’s polar ejection force activity and its inhibition during anaphase, leading to an intermediate phenotype that becomes more pronounced when endogenous KIF22 is depleted. The work shown here supports a model where the R49Q substitution partially affects KIF22 inhibition at the metaphase to anaphase transition, impeding chromosome separation to a lesser extent (**Figure 6**). This substitution could result in a population of partially inhibited KIF22 molecules that fluctuate between conformations that sequester them to microtubules (inhibited) or conformations that allow them to bind both microtubules and chromosomes (active) and generate PEF, resulting in a moderate phenotype that only becomes more severe with KIF22 knockdown. Instead of being constitutively activated like the other motor head variants, the R49Q construct is dysregulated because it exists as a mixed population of active and inactive KIF22 molecules, which would also explains the mild reduction in PEF produced by KIF22-R49Q constructs (**Figure 6**). Considering that KIF22-R49Q is a loss-of-function variant, we modeled how anaphase B segregation is affected by the loss of endogenous KIF22 function in chondrocytes via KIF22 knockdown. In these cells, loss of KIF22 activity increased the maximum distance chromosomes were able to separate. Unexpectedly, spindle pole separation was not affected by the loss of KIF22 activity, while persistent KIF22 activity from SEMDJL2 variants did impair spindle pole separation during anaphase. KIF22 activity during anaphase is incompletely understood. KIF22-WT remains on chromosomes during anaphase as seen here and in previous work (Thompson et al., 2022), and in mouse oocytes, KIF22 was shown to localize in between microtubules and anaphase chromosomes in a striated pattern and is thought to contribute to chromosome compaction in anaphase to mitigate lagging chromosomes (Oshugi et al., 2008). In the absence of KIF22 activity, we observed that chromosome separation increased, indicating that KIF22 activity on anaphase chromosomes normally provides mild resistance to anaphase B spindle pole separation. One possibility is that KIF22 activity is normally reduced during anaphase but not fully switched off. The measured activity for KIF22-T463A “inactive” phosphonull constructs does vary across cell types (Thompson et al., 2022, Soeda et al., 2016), which suggests that KIF22 activity during anaphase has different importance across cell types. However, it is mechanistically uncertain how dephosphorylated KIF22 could mediate this process, and future studies are needed to understand how KIF22 is structurally involved with chromosome organization throughout mitosis, and how this organization changes upon anaphase onset.

It is still unclear how these substitutions mechanistically interfere with KIF22 regulation at the metaphase to anaphase transition. The mechanism of KIF22 inhibition is still unknown; several possibilities have been proposed including head-to-tail autoinhibition (Thompson et al., 2022) or tail domain-mediated sequestering of KIF22 to microtubules (Soeda et al., 2016). KIF22 activity on chromosomes has been thought to be the result of collective action of many single motors rather than stable multimeric complexes like those seen for other mitotic motors like CENP-E or KIF11/Eg5 (Yajima et al., 2002). Future work remains to resolve the structure of KIF22 in its various mitotic contexts (e.g. bound to microtubules, bound to DNA, as an oligomer) to completely understand how SEMDJL2 variants mechanistically interfere with KIF22 activity.

An unexpected finding was that the severity of mitotic defects did not uniformly predict downstream nuclear architecture abnormalities. The P144T variant produced a higher percentage of abnormal nuclei, whereas E222Q caused more pronounced defects in chromosome separation. This suggests that while anaphase mis-segregation can contribute to nuclear defects (Liu et al., 2018; Hayashi et al., 2013), the relationship between the integrity of chromosome segregation and nuclear architecture is not strictly linear and are influenced by partially separable pathways. Taken together, these results highlight that anaphase dysregulation alone does not fully define how daughter cell nuclear morphology is remodeled and raises the possibility that KIF22 influences nuclear integrity through additional functions beyond anaphase mechanics.

Although the distinct mechanistic categories suggested by our studies, constitutive activation and mixed-state dysregulation, provide a more nuanced view of KIF22 function, they do not yet fully explain why mitotic defects caused by SEMDJL2 KIF22 variants selectively disrupt the development of skeletal tissue. KIF22 is expressed broadly across proliferative cell types (Uhlen et al., 2015), but the relative importance of KIF22 activity for successful mitosis seems to vary across species and cell types (Funabiki and Murray, 2000; Levesque and Compton, 2001; Tokai-Nishizumi et al., 2005; Ohsugi et al., 2008; Matthies et al., 2011). Furthermore, Dubail et al., have reported altered aggrecan secretion and extracellular matrix formation in SEMDJL2 patient-derived fibroblast cells (with the R49Q homozygous variant), and Kawaue et al., report that KIF22 KD affected chondrocyte matrix synthesis. These observations raise the possibility that impaired mitosis may intersect with chondrocyte-specific roles for KIF22 in trafficking ECM components during development, compounding the consequences of altered KIF22 activity during growth plate expansion.

Overall, this study expands the mechanistic and genotypic landscape of SEMDJL2 by identifying two novel heterozygous variants and defining how recessive and non-hotspot substitutions perturb KIF22 regulation during mitosis. The combination of clinical data and functional characterization provides compelling evidence supporting the reclassification of KIF22 p.(Pro144Thr) and p.(Glu222Gln) as likely pathogenic variants in accordance with American College of Medical Genetics and Genomics guidelines. More broadly, our findings emphasize that KIF22 function during anaphase is regulated by a finely tuned inhibitory system, disruption of which produces variant-specific consequences that likely contribute in distinct ways to skeletal dysplasia. Future structural and biochemical investigations will be essential to resolving how KIF22 transitions between its active and inhibited states, how these transitions vary across cell types, and how mitotic dysregulation intersects with chondrocyte-specific developmental programs.

## MATERIALS AND METHODS

### Cell culture and transfections

CHON-001 cells were purchased from ATCC (CRL-2846). CHON-001 cells were cultured at 37°C with 5% CO_2_ in MEM-α medium (Life Technologies, 12561072) supplemented with 20% Fetal Bovine Serum (FBS; Life Technologies, 16000044). CHON-001 acceptor cells for recombination were maintained in 5% blasticidin (ThermoFisher Scientific, R21001). CHON-001 cell lines were authenticated by STR DNA fingerprinting using the Promega GenePrint® 10 System according to manufacturer’s instructions (Promega #B9510). The STR profiles were compared to known ATCC fingerprints (ATCC.org), and to the Cell Line Integrated Molecular Authentication database (CLIMA) database version 0.1.200808 (http://bioinformatics.istge.it/clima/) (Nucleic Acids Research 37:D925-D932 PMCID: PMC2686526). The STR profiles matched known DNA fingerprints.

siRNA transfections were performed using RNAiMax (Life Technologies) according to manufacturer’s recommendations. For siRNA transfections in a 35 mm dish format, approximately 40,000 cells in 500 µL MEM-alpha medium with 10% FBS were treated with 70 pmol siRNA and 7 µL RNAiMax (Life Technologies) precomplexed for 10 minutes in 230 µL OptiMeM (Life Technologies). Cells were treated with KIF22 Silencer Validated

siRNA (Ambion #AM51331, sense sequence GCUGCUCUCUAGAGAUUGCTT). Control cells were transfected with Silencer Negative Control siRNA #2 (Ambion #AM4613) for 24 hr before fixing or imaging. Treatment volumes were adjusted to account for surface area of plates when performing siRNA KD in a 24-well format.

### Generation & validation of CHON-001 inducible lines

High-efficiency low-background (HILO) recombination mediated cassette exchange (RMCE) technology was used to engineer cell lines with one insertion of a GFP-tagged KIF22 sequence under control of a doxycycline-inducible promoter (Khandelia et al., 2011; Fonseca et al., 2019; Thompson et al., 2022; Schutt et al., 2024). To establish the RMCE acceptor cell lines, ∼60% confluent CHON cell cultures were infected with serial dilutions of the pEM584 lentiviral stock. Cells were incubated for 48 hr before the addition of selection media containing 20 μg/ml blasticidin. Next, individual clones were isolated by plating single cells via serial dilutions to a 96-well plate, and growing colonies. Clones were maintained in media with 10 μg/ml blasticidin. To generate the KIF22-GFP expressing cell lines, a RMCE donor plasmid was designed containing a siRNA-resistant KIF22-GFP construct and a puromycin resistance gene flanked by loxP sites. The KIF22-GFP donor plasmid was generated from the eGFP plasmid for recombination (pEM791) (Khandelia et al., 2011) in a previous study (Thompson et al., 2022). The KIF22-GFP plasmid was used to generate the KIF22-E222Q and P144T RMCE plasmids via QuikChange site directed mutagenesis (Agilent). The KIF22-GFP R49Q RMCE donor plasmid was generated via mutagenesis of the KIF22-GFP RMCE donor plasmid (GenScript).

The same clone of the CHON-001 acceptor cell line was used to generate all of the cell lines used in this study. Cells were electroporated using the 4D-Nucleofector® X Unit (Lonza #AAF-1003X). Approximately 1 million cells were used per reaction. Cells were spun down and resuspended in a mixture of P3 cell line Lonza Nucleofector Solution (82 µL) and Supplement Solution (18 µL). Cells were co-nucleofected with a KIF22 RMCE donor plasmid and a pEM784 nl-Cre recombinase plasmid at a 1:10 wt/wt ratio with 1350ng of total DNA in a Lonza 100 µL cuvette. After electroporation, cells sat at RT for approximately 1 min, after which cells were resuspended in 500uL of warm media, and plated in a 6-well plate, split between three wells, with each well already containing 2 mL pre-warmed MEM-alpha media supplemented with 20% FBS. 24 hr post-electroporation, media containing 5 μg/ml puromycin (Thermo Fisher Scientific, A11138-03) was added to each well to select for cells with successful recombination events. For CHON-001 lines, after 48 hr, the concentration was increased to 10 μg/ml puromycin for stronger selection. After 48 hr of strong selection, the concentration was decreased back to 5 μg/ml for maintaining the cell lines. Selected cells were pooled between the three wells. Concentrations for selection of acceptor or inducible cell lines was determined by performing a kill curve for each antibiotic. Genomic DNA from each inducible cell line was extracted (QIAmp DNA Blood Mini Kit Qiagen #51106) and sequenced through each of the variant sites (Eurofins) to verify correct incorporation of the desired KIF22 construct. Expression of KIF22-GFP constructs was induced with addition of 2 μg/ml doxycycline (Thermo Fisher Scientific #BP26531) to cell culture media. The cell lines described here are available on request from the authors.

We generated CHON-001 cells with inducible expression of GFP-tagged KIF22-WT or containing the P144T, E222Q, R49Q, or R149Q KIF22 variants as described previously (Khandelia et al., 2011; Marquis et al., 2021; Thompson et al., 2022; Queen et al., 2024; Schutt et al., 2024). Expression of constructs was induced by addition of 2 μg/ml doxycycline to cell culture media and led to an average of 1.6 to 2.5-fold higher KIF22 levels during mitosis compared to endogenous KIF22 in cells expressing GFP only (**Supplemental Figure 1A, 1B**). Importantly, this increase in expression was similar across each line (**Supplemental Figure 1B).** Additionally, these GFP-tagged KIF22 constructs contain silent mutations to confer resistance to KIF22 siRNA, enabling us to knockdown expression of endogenous KIF22 while inducing expression of KIF22-GFP simultaneously. SiRNA-mediated KIF22 depletion was quantified by immunofluorescence, resulting in 75% knockdown (**Supplemental Figure 1C, 1D**). All three of the KIF22 variant constructs localized similarly to KIF22-WT; primarily in the nucleus in interphase cells, and on chromosomes and spindle microtubules in mitotic cells (**Supplemental Figure 1A**).

### Cell fixation and immunofluorescence

Cells were seeded on acid-treated 12-mm glass coverslips in a 24-well dish and fixed in ice-cold methanol (Fisher Scientific) containing 1% paraformaldehyde (Electron Microscopy Sciences) for 10 minutes on ice and then washed 3 times for 5 minutes each in Tris-Buffered Saline (TBS; 150 mM NaCl, 50 mM Tris base, pH 7.4). Coverslips were blocked with 20% goat serum in antibody-diluting buffer (Abdil: TBS pH 7.4, 1% bovine serum albumin, 0.1% Triton-X, 0.1% sodium azide) for 1 hr at room temperature with shaking. After blocking, coverslips were washed two times for 5 minutes each in 1X TBS. Primary antibodies were diluted in Abdil: mouse anti-GFP (1:1000, 0.5 μg/ml, ThermoFisher), rat anti-tubulin YL (1:1500, 0.5 μg/ml, Millipore), rabbit anti-KIF22 (1:500, GeneTex #GTX112357). Coverslips were incubated with diluted primary antibodies overnight at 4C with shaking. Coverslips were then washed three times for 5 minutes each with 1X TBS, before incubation at room temperature with secondary antibodies against mouse, rat, and rabbit IgG conjugated to Alex Fluor 488, 594, 647 (1:500, 1 μg/ml, Molecular Probes). All secondary antibodies were diluted 1:500 in Abdil. After incubation with secondary antibodies for 1 hr at RT with shaking, coverslips were washed 2 times in 1X TBS for 5 minutes each, and 1 time in 1X TBS with 0.125 μg/mL DAPI, and then washed 1 time in 1X TBS before mounting in Prolong Gold anti-fade mounting medium (Invitrogen Molecular Probes, P36935).

### KIF22 expression level quantification in CHON-001 inducible lines

The efficiency of siRNA-mediated KIF22 depletion in CHON-001 cells was measured via quantitative fluorescence measurements of KIF22 antibody signal on mitotic spindles in ImageJ. Cells from the CHON GFP-only line were treated with control or KIF22 siRNA for 24 hr and then fixed and stained for KIF22 in a 24-well format as described above. Images were acquired from a minimum of 3 experimental replicates. Images were background subtracted, and intensity was measured using a mask of the tubulin signal for each mitotic cell to quantify KIF22 signal. Fluorescence intensities were normalized to the average intensity of the control siRNA treated condition. (**Supplemental Figure 1**).

The expression of KIF22-GFP tagged constructs was also measured via quantitative fluorescence measurements of KIF22 antibody signal on mitotic spindles in ImageJ. Cells from each inducible chondrocyte line used here were plated in a 24-well plate on glass coverslips. 24 hr after plating, the media was exchanged to media containing 2 μg/ml doxycycline to induce expression of GFP-tagged constructs. 6 hr later, cells were fixed and stained as described above for KIF22, and tubulin. KIF22 intensity in mitotic cells was measured in ImageJ as described for the siRNA efficiency measurements, and compared across lines. Fluorescence intensities were normalized to that of the GFP-only chondrocyte line, which expresses only endogenous KIF22 (**Supplemental Figure 1**).

### Live cell imaging of anaphase chromosome segregation & analysis

For live cell imaging of chromosome segregation in anaphase chondrocyte spindles, approximately 40,000 CHON-001 cells were seeded in 35mm glass bottom plates and left to settle overnight. For experiments where endogenous KIF22 was depleted, cells were treated with siRNA as described above for 24 hr. After siRNA treatment, the media on the plates was exchanged to CO2-independent media containing 2 μg/ml doxycycline and 100 nM SiR Tubulin (Spirochrome #SC002) and supplemented with 20% FBS approximately 2 hr before imaging. Movies were taken of mitotic cells in metaphase with 3 focal plane images acquired at 1-min time intervals using a 60X objective until cytokinesis was completed. Analysis of chromosome segregation and spindle pole separation was performed as described previously (Thompson et al., 2022). Distances between segregating chromosome masses were measured by plotting the GFP intensity along a line drawn through both spindle poles (macro available at https://github.com/StumpffLab/Image-Analysis; Stumpff, 2021). This data set was split at the center distance to generate two plots, each representing one half-spindle/segregating chromosome mass. Next, the distance between the maximum of each intensity plot was calculated using MATLAB (MathWorks, version R2018a) (script available at https://github.com/StumpffLab/Image-Analysis).

### Nuclear morphology analysis in fixed cells

CHON-001 cells were seeded in 24-well plates and fixed as described above. Cells were treated with 2 μg/ml doxycycline 16 hours before fixation to induce expression of GFP constructs. Cells were fixed as described above. Nuclei of interphase cells were imaged at 100X. Random fields of view were captured, and approximately 100 cells per condition, per experiment were analyzed. Nuclear solidity was measured in ImageJ, and the fifth percentile of solidity in GFP-only control cells was used as a threshold to sort nuclei into “normal” or “abnormal” shapes.

### Polar ejection force analysis in fixed cells

CHON-001 cells were seeded in 24-well plates and fixed as described above. Cells were treated with 2 μg/ml doxycycline 16 hours before fixation to induce expression of GFP constructs. Cells were treated with 100 μM monastrol (Selleckchem #S8439) for 2–3 hr before fixation. Monopolar mitotic cells were imaged using a 100X objective such that the spindle poles were in focus. Polar ejection forces were measured as described previously (Thompson et al., 2022). Briefly, a circular ROI with a radius of 14 μm was centered around the spindle poles of each monopolar spindle. Next, the radial profile of the DAPI signal intensity was measured using the Radial Profile Plot plugin in ImageJ (https://imagej.nih.gov/ij/plugins/radial-profile.html). The distance from the pole to the maximum DAPI signal intensity was determined for each cell as a proxy for relative polar ejection forces.

### Microscopy

Fixed cell images and live cell movies were acquired on Ti-2E inverted microscopes (Nikon Instruments) driven by NIS Elements (Nikon Instruments). Images were captured using either a Photometrics sCMOS Prime BSi camera, or an Andor Zyla sCMOS camera. Imaging was performed using the following Nikon objectives: Plan Apo λ 60 × 1.42 NA, and APO 100 × 1.49 NA. For live cell-imaging, cells were imaged in CO_2_-independent media (Life Technologies #18045-088) supplemented with 20% FBS (Life Technologies #16000-044) within environmental chambers held at 37°C.

---

**Supplemental Figure 1:**
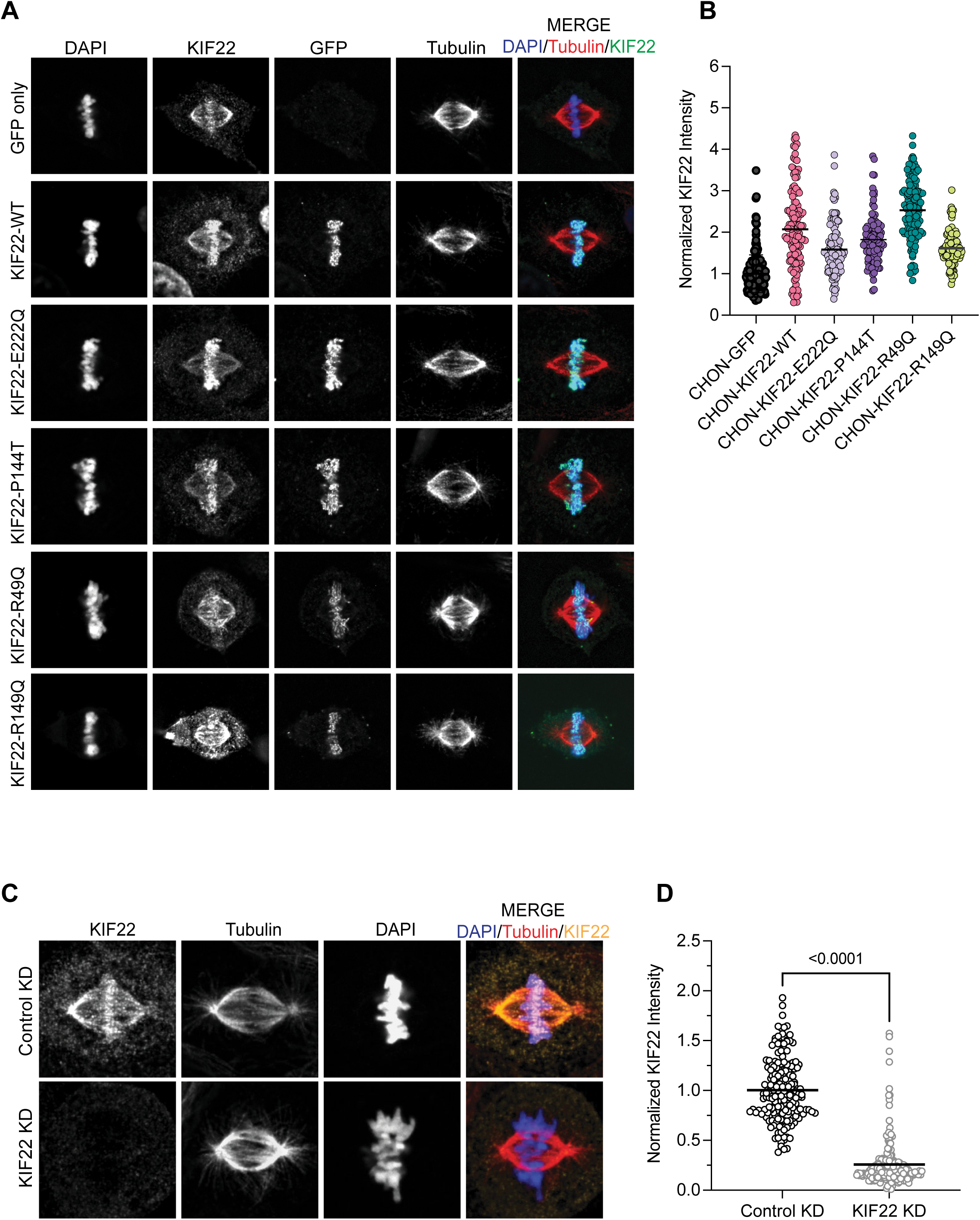
Chondrocyte cell line validation. **(A)** Representative images of fixed chondrocytes from each line used in this study. DNA, KIF22, GFP, and tubulin were labeled in each line. **(B)** KIF22 intensity was measured in mitotic cells from each chondrocyte cell line. Intensity values were normalized to the average KIF22 intensity observed in the GFP-only chondrocyte line. Cell counts – GFP: 101 cells; KIF22-WT: 126 cells; KIF22-E222Q: 120 cells; KIF22-P144T: 104 cells. **(C)** Representative images of fixed GFP-only expressing chondrocytes treated with either control or KIF22 siRNA, and with KIF22, DNA, and tubulin labeled. (D) Quantification of KIF22 intensity in mitotic cells treated with either control siRNA or KIF22 siRNA. Data were compared by running an unpaired t-test. Cell count – control KD: 178 cells; KIF22 KD: 186 cells.

