## Supplemental Figures for "Novel KIF22 Variants Disrupt Mitosis in Human Chondrocytes and Expand SEMDJL2 Mechanisms"

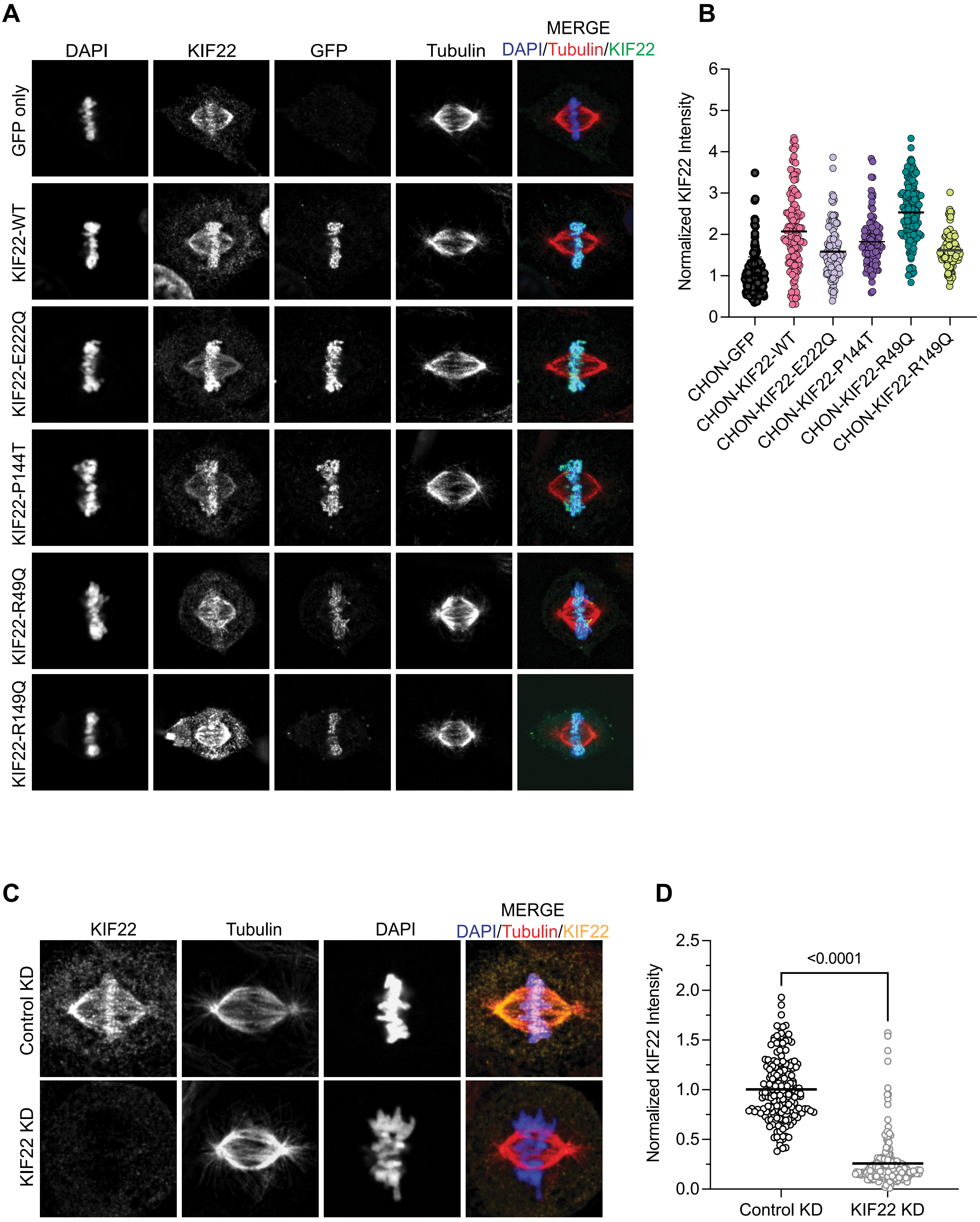
A
DAPI
KIF22
GFP
Tubulin
MERGE
DAPI/Tubulin/KIF22
GFP only
KIF22-WT
KIF22-E222Q
KIF22-P144T
KIF22-R49Q
KIF22-R149Q
B
Normalized KIF22 Intensity
0
1
2
3
4
5
6
CHON-GFP
CHON-KIF22-WT
CHON-KIF22-E222Q
CHON-KIF22-P144T
CHON-KIF22-R49Q
CHON-KIF22-R149Q
C
KIF22
Tubulin
DAPI
MERGE
DAPI/Tubulin/KIF22
Control KD
KIF22 KD
D
Normalized KIF22 Intensity
0.0
0.5
1.0
1.5
2.0
2.5
<0.0001
Control KD
KIF22 KD

**Supplemental Figure 1: Chondrocyte cell line validation. (A)** Representative images of fixed chondrocytes from each line used in this study. DNA, KIF22, GFP, and tubulin were labeled in each line. **(B)** KIF22 intensity was measured in mitotic cells from each chondrocyte cell line. Intensity values were normalized to the average KIF22 intensity observed in the GFP-only chondrocyte line. Cell counts – GFP: 101 cells; KIF22-WT: 126 cells; KIF22-E222Q: 120 cells; KIF22-P144T: 104 cells. **(C)** Representative images of fixed GFP-only expressing chondrocytes treated with either control or KIF22 siRNA, and with KIF22, DNA, and tubulin labeled. (D) Quantification of KIF22 intensity in mitotic cells treated with either control siRNA or KIF22 siRNA. Data were compared by running an unpaired t-test. Cell count – control KD: 178 cells; KIF22 KD: 186 cells.
